# Pollutants in Hong Kong Soils: As, Cd, Cr, Cu, Hg, Pb and Zn

**DOI:** 10.1101/2020.02.16.951558

**Authors:** M.K. Chung, K.C. Cheung, M.H. Wong

**Affiliations:** Croucher Institute for Environmental Sciences and Department of Biology, Hong Kong Baptist University, Kowloon Tong, Kowloon, Hong Kong SAR, PR China

**Keywords:** Urban soils, South China, Heavy Metals, Contamination, Pollution

## Abstract

Six heavy metals (Hg, Cu, Cd, Cr, Pb, Zn) and 1 metalloid (As) in surface soils of Hong Kong were investigated in 10 land use categories (urban park, greening area, country park, rural area, restored landfill, agricultural farmland, orchard farm, crematorium, industrial and near highway area). Edaphic Hg concentration in Hong Kong was firstly reported here. Clustering of land uses was observed based on total pollutants concentrations (sum of 7 metals). The most polluted cluster consisted of industrial and highway areas (median: 617 to 833 mg kg^-1^) and the runner-up cluster included urban park, greening area and restored landfill (median: 400 to 500 mg kg^-1^). However, this general finding was not observed for Hg, where higher concentration was found in agricultural farmland (median 109 μg kg^-1^). The use of low quality fertilizers, together with the contribution from exhausts and wearable parts from automobiles were believed to be the major sources of Cr, Cu and Zn in Hong Kong, while the application of Hg-containing agrochemicals maybe the main mechanism of Hg contamination in agricultural soil. Based on the daily intake assumption of 0.2 g d^-1^ of soil particles by USEPA, direct ingestion of Hg-containing soils is not a major exposure pathway for population in Hong Kong. When comparing the edaphic heavy metal concentrations with Dutch soil quality guidelines demonstrated that Hg, Cd and Pb were not in level of health concerns, while Cu, Cr and Zn in less than 6% of total samples were found to exceed the Dutch intervention values sporadically. In contrast, suburban soils from northern and northeastern Hong Kong were mostly contaminated with As (10% of total samples) at concentration that could be potentially causing adverse health impacts to the nearby population.

**Pollutant in Hong Kong soils series:**

1. Chung, M. K., Hu, R., Cheung, K. C. & Wong, M. H. Pollutants in Hong Kong soils: Polycyclic aromatic hydrocarbons. Chemosphere 67, 464–473 (2007). http://doi.org/10.1016/j.chemosphere.2006.09.062
2. Chung, M. K., Cheung, K. C. & Wong, M. H. Pollutants in Hong Kong Soil: As, Cd, Cr, Cu, Hg, Pb and Zn https://doi.org/10.1101/2020.02.16.951558
3. Chung, M. K., Hu, R., Cheung, K. C. & Wong, M. H. Pollutants in Hong Kong Soils: Organochlorine Pesticides and Polychlorinated Biphenyls https://doi.org/10.1101/2020.02.16.951541

## 1. Introduction

Environmental contamination of heavy metals has been reported world-wide, and excessive exposure to toxic metals has been commonly known to be hazardous to human health (Agency for Toxic Substances and Disease Registry, 2006). Among these toxic metals, Hg is of particular in concern as it is characterized by high vapour pressure. This unique feature makes it ubiquitous in the environment and become a global pollutant. In addition, its toxicity to human, especially childbearing women, also highlighted the concern from government agency to work on Hg reduction (Srivastava *et al.*, 2006). In countries under the European Union, Hg is classified as a dangerous chemical because of its mobility, volatility and its bioaccumulative properties within organisms and along the food chains (Mukherjee *et al.*, 2004).

A recent review on Hg contamination in China (Zhang and Wong, 2006) revealed that soil Hg contents in most cities (70 to 700 μg kg^-1^, 12 out of 14 cities reviewed) exceeded the background edaphic Hg value in China (65 μg kg^-1^) (State Environmental Protection Administration of China, 1990). Enrichment of Hg and other metals was observed in agricultural crop soils (Wong *et al.*, 2002) and sediments (Cheung *et al.*, 2003) around the Pearl River Delta. This led to higher concentrations of all the toxic metals found in bivalves and freshwater fish collected (including from the field and available in markets) within the region (Fang *et al.*, 2001, 2003; Kong *et al.*, 2005; Zhou and Wong, 2000), including seafood in Hong Kong (Tam and Mok, 1991).

In Hong Kong, it has been observed that Hg was bioaccumulated in cetaceans and Indo-Pacific hump-backed dolphins (*Sousa chinensis*) (Parsons, 1998, 1999). It was noted that the mean value of Hg in adult human hair was 3.3 μg g^-1^ which was higher the mean value for US counterpart (1.5 μg g^-1^) (Dickman and Leung, 1998), also the recommended limit for Hg in hair set by USEPA is 1 μg g^-1^ (Gallagher, 2006). It was indicated that the elevated Hg levels were linked to subfertility in Hong Kong males (Dickman *et al.*, 1998). In addition, higher Hg levels in blood and hair of children in Hong Kong were also observed to be correlated with the frequency of fish consumption (Ip *et al.*, 2004).

Soils can act as both sinks, via dry atmospheric deposition; as well sources of tosic metals, via re-emission of semi-volatile pollutants and wind-blown of contaminated soil materials. By direct contact and inhalation of soil particles, toxic metals would pose risks to human health. In addition, leaching is also one of the important pathways to transfer toxic metals to water bodies and therefore accumulate in various aquatic organisms. Ultimately the toxic metals can enter human via consumption of food such as crops and fish, and thus understanding the levels and potential sources of toxic metals are essential to secure public health. Modeling of exposure mechanisms such as dermal contact and inhalation of dust of soil pollutants for risk assessment requires intensive data (U.S. Environmental Protection Agency, 1996c), and there is a lack of data on toxic metals especially Hg in Hong Kong soils. A more comprehensive survey of toxic metals (except Hg) in Hong Kong soils was conducted almost 10 years ago (Chen *et al.*, 1997), and therefore there seems to be a need to provide update information for all the toxic metals contained in soils.

The present study was aimed to address the concerns mentioned above by providing current status of heavy metal and metalloid concentrations (Hg, As, Cu, Cd, Cr, Pb and Zn) in Hong Kong soils, with a special focus on Hg. To our knowledge, this is the first report on edaphic Hg levels in Hong Kong and the nearby Pearl River Delta. Results are valuable as they partially filled the lacking edaphic metals (especially Hg) information in the Delta and can act as reference for other studies in the region where contamination of Hg is in raising concern by the public. Potential sources of these toxic metals are also discussed.

## 2. Materials and Methods

### 1. Sampling and Analysis

The sampling was based on 10 different land uses in Hong Kong: urban park, country park, rural area, restored landfill, agricultural farmland, orchard farm, crematorium, industrial area and nearby highway. All together there were 138 composite soil samples that taken from the depth of 0 to 5 cm from surface by a stainless steel soil core. Samples were stored in plastic bags and subsequently air-dried for 2 weeks and sieved through a 2-mm mesh.

Chemical analyses of metal contents in soils were based on standard method. 0.25 g of soil sample was mixed with 9 mL nitric acid, 3 mL hydrofluoric acid and 1 mL hydrochloric acid and subjected to microwave-assisted acid digestion (USEPA 3052) (U.S. Environmental Protection Agency, 1996a). The solutions were then filtered through Advantec 5C filter paper, diluted and made up with deionized water in a 50-ml plastic volumetric flask. Concentrations of As, Cu, Cd, Cr, Pb, and Zn were determined by inductively coupled plasma - optical emission spectrometry (ICP-OES) (Perkin-Elmer Optima 3000 DV), while Hg was quantified by Flow Injection Mercury System (FIMS) (Perkin-Elmer FIMS-400) based on the cold-vapor atomic absorption spectrometry (CVAA) (U.S. Environmental Protection Agency, 1996b). Limit of detection (LOD) for Hg was 0.5 μg kg^-1^, while 50 μg kg^-1^ for Cd, Cr, Cu and Zn, and 100 μg kg^-1^ for As and Pb.

### 2. Quality assurance and Data analysis

Standard Reference Material (SRM) 2711 was obtained from National Institute of Standards and Technology (NIST, USA). An analytical blank and the SRM were included in every batch of microwave acid digestion to assess the recoveries and performance of extraction.

Mean individual recoveries were: 83 ± 2% (Hg), 102 ± 5% (As), 88 ± 1% (Cd), 97 ± 1% (Cu), 107 ± 4% (Pb) and 89 ± 2% (Zn). On average, the recoveries of all the investigated elements in SRM were all > 94%. Statistical analyses including descriptive statistics, correlation analysis, and PCA analysis were conducted with Statistica (version 6.0 from StatSoft). Not detected values were substituted with half of lowest limit of detection (LOD) only for descriptive statistics.

## 3. Results and Discussion

### 1. Concentration of pollutants in Hong Kong

Kriged maps were constructed to show the spatial distribution of investigated pollutants (Figure 1), and it is observed that most of the hotspots for pollutants were found in the northern part of Hong Kong. In addition, clustering of soil pollutant concentrations in land uses was also observed (Table 1). Total heavy metal concentrations were highest in industrial area and area nearby highway (median 617 and 833 mg kg^-1^); while similar for urban park, greening area and restored landfill (median 400 to 500 mg kg^-1^), and the rest of the land uses are least contaminated (median 200 to 350 mg kg^-1^). The general findings that soils in industrial area and adjacent to highways were most contaminated can also be observed when considering the pollutant individually, but excluding Hg. Variations in pollutant concentration were usually greatest in urban park, which spanned up to 3 orders of magnitude, and large variations were found in most of the land uses, which reflects the heterogeneity of pollutants concentrations is under the strong influence of local activities or pollution sources.

**Figure 1.**
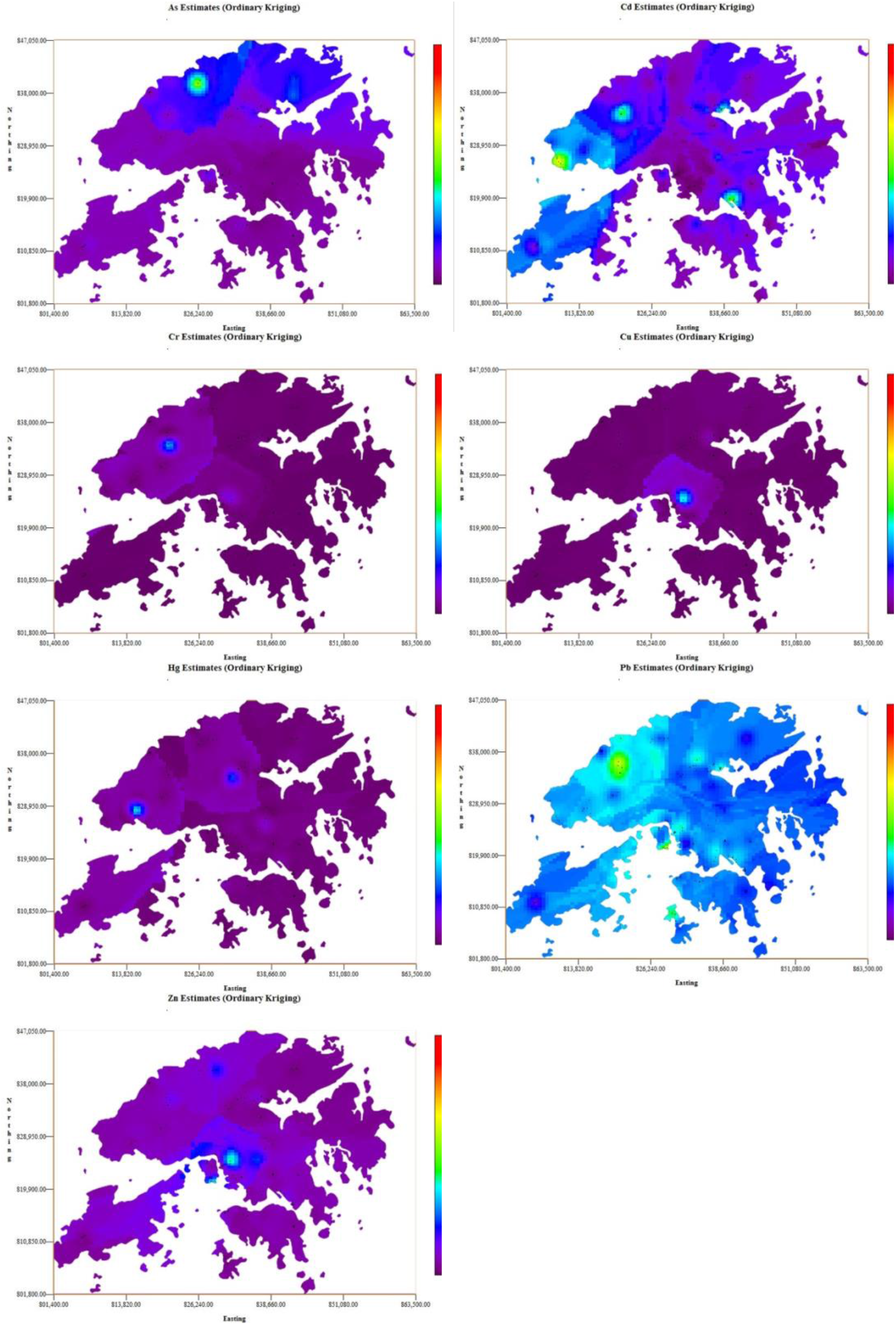
Kriged maps of pollutants (As, Cd, Cr, Cu, Hg, Pb, and Zn) concentrations (mg kg^-1^) in surface soils of Hong Kong.

**Table 1.** Mean, median, and range of concentration of studied pollutants (mg kg^-1^) in Hong Kong soils. Note that concentration unit of Hg μg kg^-1^.

| Classified soil categories | Sample no. | Hg |  |  | As |  |  | Cu |  |  | Cd |  |  |
| --- | --- | --- | --- | --- | --- | --- | --- | --- | --- | --- | --- | --- | --- |
|  |  | Mean | Median | Range | Mean | Median | Range | Mean | Median | Range | Mean | Median | Range |
| | | Concentration in soils ($\mu\text{g/kg}$) | | | Concentration in soils ($\text{mg/kg}$) | | | Concentration in soils ($\text{mg/kg}$) | | | Concentration in soils ($\text{mg/kg}$) | | |
| Urban park | 39 | 232 | 75.0 | [N.D. - 3785] | 19.4 | 16.5 | [N.D. - 122] | 81.41 | 13.0 | [N.D. - 2129] | 0.56 | 0.01 | [N.D. - 2.93] |
| Greening area | 14 | 104 | 66.0 | [N.D. - 311] | 26.6 | 26.6 | [N.D. - 97] | 21.46 | 15.1 | [N.D. - 59] | 0.50 | 0.01 | [N.D. - 2.25] |
| Country park | 9 | 52 | 25.0 | [N.D. - 128] | 36.1 | 20.3 | [3.6 - 95] | 11.0 | 0.01 | [N.D. - 93] | 0.36 | 0.01 | [N.D. - 2.42] |
| Rural area | 19 | 75 | 81.0 | [N.D. - 202] | 27.6 | 20.5 | [3.2 - 143] | 7.32 | 6.08 | [N.D. - 28] | 0.37 | 0.01 | [N.D. - 2.91] |
| Restored landfill | 11 | 50 | 25.0 | [N.D. - 119] | 22.0 | 22.8 | [N.D. - 47] | 19.3 | 11.0 | [N.D. - 62] | 0.85 | 0.01 | [N.D. - 2.86] |
| Agricultural farmland | 9 | 402 | 109 | [N.D. - 2196] | 27.2 | 15.2 | [N.D. - 109] | 12.0 | 6.61 | [N.D. - 66] | 0.14 | 0.01 | [N.D. - 0.77] |
| Orchard farm | 5 | 154 | 72.0 | [N.D. - 555] | 34.3 | 30.4 | [7.3 - 93] | 8.86 | 5.60 | [N.D. - 27] | 0.56 | 0.01 | [N.D. - 1.6] |
| Crematorium | 10 | 75 | 78.0 | [N.D. - 146] | 13.8 | 11.5 | [N.D. - 50] | 4.29 | 0.61 | [N.D. - 16] | 0.13 | 0.01 | [N.D. - 1.21] |
| Industrial area | 18 | 74 | 73.0 | [N.D. - 237] | 25.5 | 25.4 | [N.D. - 61] | 52.34 | 33.3 | [N.D. - 396] | 1.19 | 0.67 | [N.D. - 4.11] |
| Nearby highway | 4 | 134 | 117 | [N.D. - 277] | 174.6 | 141 | [80.5 - 336] | 18.75 | 15.4 | [3.62 - 41] | 0.01 | 0.01 | [N.D. - N.D.] |

| Classified soil categories | Sample no. | Cr |  |  | Pb |  |  | Zn |  |  | Total pollutants |  |  |
| --- | --- | --- | --- | --- | --- | --- | --- | --- | --- | --- | --- | --- | --- |
|  |  | Mean | Median | Range | Mean | Median | Range | Mean | Median | Range | Mean | Median | Range |
| | | Concentration in soils ($\text{mg/kg}$) | | | | | | | | | | | |
| Urban park | 39 | 50.5 | 21.9 | [1.85 - 601] | 141 | 130 | [11 - 305] | 293 | 197 | [13 - 3508] | 586 | 401 | [29 - 6553] |
| Greening area | 14 | 20.8 | 18.2 | [1.23 - 44] | 142 | 144 | [104 - 188] | 367 | 300 | [114 - 1182] | 578 | 540 | [300 - 1351] |
| Country park | 9 | 7.18 | 2.00 | [N.D. - 23] | 75 | 68 | [56 - 115] | 75 | 85 | [38 - 110] | 204 | 204 | [101 - 357] |
| Rural area | 19 | 17.0 | 12.1 | [0.81 - 57] | 136 | 124 | [53 - 244] | 183 | 135 | [43 - 555] | 371 | 377 | [155 - 763] |
| Restored landfill | 11 | 25.1 | 24.9 | [11.8 - 37] | 189 | 180 | [47 - 393] | 170 | 146 | [79 - 347] | 426 | 425 | [221 - 680] |
| Agricultural farmland | 9 | 22.7 | 17.9 | [7.36 - 54] | 121 | 120 | [79 - 161] | 197 | 152 | [63 - 564] | 380 | 303 | [234 - 845] |
| Orchard farm | 5 | 16.3 | 12.4 | [5.73 - 34] | 104 | 91 | [64 - 186] | 116 | 95 | [45 - 243] | 280 | 236 | [145 - 479] |
| Crematorium | 10 | 27.3 | 21.1 | [11.5 - 88] | 118 | 96 | [65 - 277] | 140 | 126 | [89 - 221] | 304 | 263 | [180 - 515] |
| Industrial area | 18 | 167 | 29.1 | [11.3 - 2486] | 221 | 183 | [92 - 493] | 529 | 298 | [75 - 1631] | 996 | 617 | [214 - 3033] |
| Nearby highway | 4 | 35.5 | 35.4 | [15.7 - 55] | 147 | 134 | [115 - 205] | 556 | 349 | [158 - 1367] | 932 | 833 | [409 - 1653] |

Hg concentration was ranged from non detectable (N.D.) to 3790 μg kg^-1^, and the 10 most contaminated soil samples were found in urban parks, greening areas and farms (29 and 3790 μg kg^-1^). The 5 locations with highest Hg levels were: urban parks in Kwun Tong (633 μg kg^-1^), Central (985 μg kg^-1^) and Tuen Mun (3785 μg kg^-1^) and agricultural farm in Sha Tin (762 μg kg^-1^) and Tai Po (2196 μg kg^-1^). The usual contents of Hg in soils are in the range of 0.01 to 0.03 μg kg^-1^ (Senesi *et al.*, 1999). For contaminated areas such as Hg mine, Hg concentrations in soils were a thousand folds more (Loredo *et al.*, 1999). Median Hg levels in urban soil in Korea and Norway were 45 and 130 μg kg^-1^ respectively (Kim and Kim, 1999; Reimann and Caritat, 1998). The mean and median Hg concentrations in Hong Kong were 135 and 70.5 μg kg^-1^ respectively, which were broadly in line with the Hg concentrations observed in major cities in China (Beijing: 509 μg kg^-1^, Chongqing: 319 μg kg^-1^, Wuhan: 314 μg kg^-1^) (Liu *et al.*, 1998; Wang, 2001; Wang *et al.*, 2005). In addition, the concentration ranges of Cd (N.D. to 4.11 mg kg^-1^), Cr (N.D. to 2500 mg kg^-1^), and Pb (11 to 490 mg kg^-1^) in the present study (Table 1) were similar to those reported in Shenyang, Beijing, Nanjing and Xi’an, China (Fang *et al.*, 2004; Wang *et al.*, 2001). However, the range of As (N.D. to 336 mg kg^-1^) was generally higher for an order of magnitude when compared with those reported in major cities in China (Wang *et al.*, 2001). This implied that there are significant sources of As in Hong Kong that are absent from the aforementioned cities. Mean concentrations of Cu (37.2 mg kg^-1^) and Zn (276 mg kg^-1^) in Hong Kong were closed to those found in Nanjing (Cu: 40.4 mg kg^-1^, Zn: 280 mg kg^-1^) (Wu *et al.*, 2003), but higher than those reported in Guangzhou (Cu: 9.62 mg kg^-1^, Zn: 115.4 mg kg^-1^) (Guan *et al.*, 2001).

### 2. Statistical analyses among pollutants and their potential sources

The correlations among Cu, Cr and Zn were also identified by principal factor 1 (PC 1) in the PCA plots shown in Figure 2a. Cadmium was excluded for PCA because of a large set of not-detected value. PC 1 was able to explain 56% of the variance while PC 2 explained 16%. Together they extracted 72% of the total variance from the present study. However, PC2 represented an antagonistic relationship between Hg and As. Figure 2b shows the projection of sampling points to the factor plane. Samples from 2 different urban parks in New Territories contained very high level of metals (Cu, Cr and Zn) and Hg. Certain agricultural farms and soils from rural and adjacent to highway in the New Territories were best explained by PC2, implied that they are either high in Hg or As concentration. Arsenic level was reported to be higher in industrial and heavy traffic sites (Deb *et al.*, 2002), and the present study also indicated higher level of edaphic As in the vicinity of highways.

**Figure 2.**
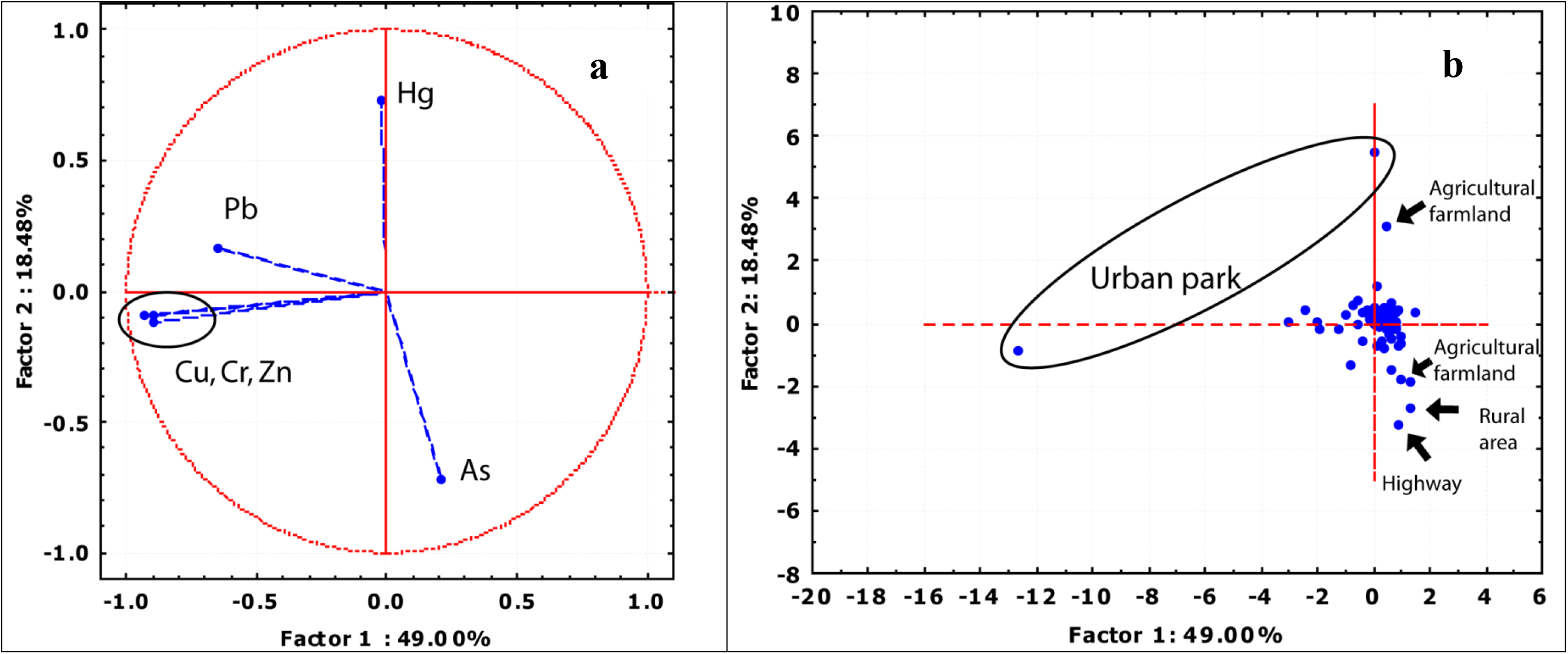
Plots with PC 1 and PC 2 from principal component analysis as X and Y axis on various pollutants and sampling points. PC 1 was able to explain 49% while PC 2 accounted for another 19% of the total variance. Note that Cd were excluded from PCA because of a large set of not-detected value. A: Biplot showing the loading of 6 pollutants on PC 1 and PC 2. B: Scatter plot of sampling points projecting on the PC 1 and PC 2 plane.

Arsenic has both natural and anthropogenic sources, and their anthropogenic origins included agrochemicals such as herbicides and pesticides (in form of monosodium methanearsonate), and wood preservative (arsenic trioxide) (USGS, 2006). Since high levels of As (>100 mg kg^-1^) were found chiefly in soil samples collected nearby highways, it is expected that automobiles could be the major contributor of As through burning of fossil fuel and wearing of the As-containing babbitt bearings. Many of the soil samples with As ranged from 30 to 100 mg kg^-1^ were collected in the northern rural part of Hong Kong, in which historical use of agrochemicals containing As would be the main contributor of As in soils.

Higher Hg levels were observed in soils of parks and farms with plantation (Table 1). Similar to As, the origin of Hg can be both anthropogenic and natural, such as ore mining and forest fire. Certain fertilizers, pesticides and fungicides are known to contain Hg (Matthews *et al.*, 1995; Nakagawa and Hiromoto, 1997). Therefore, the Hg concentrations found in farmlands, orchard farms and urban parks maybe due to the use of agrochemicals. The world-wide average of Hg content in coal is 0.1 ± 0.01 mg kg^-1^, whereas coals from southern China were enriched in Hg by 1 to 2 orders of magnitude (Yudovich and Ketris, 2005). Due to the fact that Hong Kong is located at the southern tip of PRD, which is known for its electrical and electronic manufacturing industry, the high power demand and the associated emission of Hg is likely to create a regional Hg problem (Wang *et al.*, 2006), and hence contributes to Hg level in Hong Kong soils.

Higher levels of Hg in human hair leading to subfertility in males have been suspected to link with higher rates of fish consumption in Hong Kong (Dickman *et al.*, 1998; Dickman and Leung, 1998). The average consumption rate of seafood is about 60 kg yr^-1^ person^-1^, which is equivalent to 167 g d^-1^ person^-1^, and the mean Hg levels in marine and freshwater fish available in markets were 120 and 80 μg kg^-1^ respectively (Dickman and Leung, 1998). The present study showed that the average Hg level of 135 μg kg^-1^ in Hong Kong soils is slightly higher than those reported in fish. However, the assumed ingestion amount of soil particles for children (15 kg in weight) is 0.2 g d^-1^ when calculating the soil screening level for residential exposure of soil pollutants (U.S. Environmental Protection Agency, 1996c), which is 833 folds less than the average intake rate of fish (167 g d^-1^). It is therefore believed that the direct ingestion of Hg-contaminated soils is not a major health concern in Hong Kong.

Table 2 shows the most prominent correlations were Cu-Cr, Cu-Pb and Cu-Zn, which were found in 5 out of 10 different land uses. Soil samples collected adjacent to highway, country park, agricultural farmland, crematorium and industrial area did not show more than 2 significant correlations. It is a common practice to use compost as soil conditioner in urbanized areas. In Hong Kong, the sources for composting are largely derived from livestock wastes from pig and poultry farms under the free livestock waste collection service provided by the government. Pig manure contains high levels of Cu and Zn (Bowland, 1990) as common additives in pig feed to increase the feed conversion efficiency and economic returns (Jin *et al.*, 1995). Cases of excessive addition of Cu and Zn in feeds were noted (Kessler *et al.*, 1994), and considerable amounts of these metals were also reported in local composts (Wong, 1990). Apart from compost, fertilizers are also known to contain As, Cd, Cr, Pb and Zn (Guan *et al.*, 2001; Renner, 2004). Land application of sewage sludge was also reported to be the principle sources of heavy metals, especially Cd and As (Chu and Wong, 1984; Elinder, 1985), but this possibility can be ruled out since sludge is commonly dumped in domestic landfills in Hong Kong. In England and Wales soil, greatest inputs of Zn and Cu were from animal manure and greatest inputs of Cr were from industrial wastes (McGrath, 2000). The significant correlations of Cu-Cr, Cu-Pb and Cu-Zn implied that they are derived from the same sources, and the most likely source of these metals in Hong Kong soil is from low quality fertilizers, because of its ease of application and more stable quality than compost. In addition, Zn and Cu are pollutants associated with automobiles (Viklander, 1997). Approximately 3% of ZnO is commonly added to the tyres of vehicles as a vulcanization agent and the wear of tyres can be a significant source of Zn in urban areas (Friedlander, 1973). Other heavy metal compounds including Cu, Cd and Pb (~0.002%, <0.001% and <0.005% respectively) are identified in tyres (UNEP, 2000). Other wearable parts of vehicles such as brake and brake lining also contained high contents of Cu, Pb and Zn (80 to 24 000 mg kg^-1^) (Westerlund, 2001) and therefore contribute a significant portion of heavy metals in soils.

**Table 2.** Correlation matrix of investigated metals in different land uses in Hong Kong.

| Urban park |  |  |  |  |  |  |  | Agricultural farmland |  |  |  |  |  |  |  |  |  |
| --- | --- | --- | --- | --- | --- | --- | --- | --- | --- | --- | --- | --- | --- | --- | --- | --- | --- |
|  |  | Hg | As | Cu | Cd | Cr | Pb | Zn |  |  | Hg | As | Cu | Cd | Cr | Pb | Zn |
| Greening area | Hg |  | -0.06 | 0.04 | 0.2 | 0.01 | 0.12 | 0.03 | Orchard farm | Hg |  | -0.2 | 0.16 | -0.23 | 0.13 | 0.25 | 0.08 |
|  | As | 0.46 |  | -0.04 | 0.12 | -0.07 | -0.16 | -0.76 |  | As | -0.61 |  | -0.15 | 0.36 | -0.45 | -0.55 | -0.43 |
|  | Cu | 0.13 | 0.54* |  | -0.09 | 0.87* | 0.47* | 0.98* |  | Cu | -0.19 | 0.83 |  | -0.21 | 0.62 | 0.53 | 0.86* |
|  | Cd | 0.14 | 0.47 | 0.53* |  | -0.17 | 0.15 | -0.08 |  | Cd | 0.4 | 0.84 | 0.73 |  | -0.2 | -0.65 | -0.18 |
|  | Cr | 0.14 | 0.46 | 0.80* | 0.26 |  | 0.49* | 0.82* |  | Cr | -0.14 | 0.15 | 0.66 | 0.15 |  | 0.48 | 0.90* |
|  | Pb | -0.4 | -0.16 | 0.53 | 0.19 | 0.35 |  | 0.54* |  | Pb | -0.1 | -0.35 | 0.23 | -0.27 | 0.87 |  | 0.6 |
|  | Zn | -0.08 | -0.2 | 0.1 | -0.23 | 0.33 | 0.13 |  |  | Zn | -0.17 | 0.82 | 0.99* | 0.74 | 0.62 | 0.2 |  |
| Country park |  |  |  |  |  |  |  | Crematorium |  |  |  |  |  |  |  |  |  |
|  |  | Hg | As | Cu | Cd | Cr | Pb | Zn |  |  | Hg | As | Cu | Cd | Cr | Pb | Zn |
| Rural area | Hg |  | -0.32 | -0.21 | -0.51 | 0.67* | 0.52 | 0.18 | Industrial area | Hg |  | -0.08 | 0.25 | 0.18 | 0.33 | -0.16 | 0.07 |
|  | As | 0.31 |  | 0.3 | 0.44 | 0.03 | 0.09 | 0.18 |  | As | 0.05 |  | 0.09 | 0.85* | 0.1 | 0.14 | -0.04 |
|  | Cu | 0.41 | 0.49* |  | 0.01 | 0.43 | 0.51 | 0.1 |  | Cu | 0.78* | -0.08 |  | -0.18 | 0.82* | 0.13 | 0.72* |
|  | Cd | 0.02 | 0.70* | 0.43 |  | -0.23 | -0.31 | 0.47 |  | Cd | 0.17 | 0.08 | -0.19 |  | -0.05 | -0.16 | -0.35 |
|  | Cr | 0.37 | 0.4 | 0.27 | -0.03 |  | 0.79* | 0.6 |  | Cr | 0.07 | -0.42 | 0.02 | 0.22 |  | -0.05 | 0.53 |
|  | Pb | -0.16 | -0.22 | -0.08 | -0.28 | -0.1 |  | 0.29 |  | Pb | -0.26 | -0.14 | 0.14 | -0.11 | -0.15 |  | 0.61 |
|  | Zn | 0.48* | -0.03 | 0.12 | -0.2 | 0.27 | 0.14 |  |  | Zn | 0.2 | -0.31 | 0.43 | -0.21 | -0.06 | 0.4 |  |
| Restored landfill |  |  |  |  |  |  |  |  |  |  |  |  |  |  |  |  |  |
|  |  | Hg | As | Cu | Cd | Cr | Pb | Zn |  |  |  |  |  |  |  |  |  |
| Nearby highway | Hg |  | 0.07 | -0.36 | -0.62* | -0.21 | -0.23 | 0.39 |  |  |  |  |  |  |  |  |  |
|  | As | -0.76 |  | -0.43 | -0.09 | -0.79* | -0.54 | -0.4 |  |  |  |  |  |  |  |  |  |
|  | Cu | -0.68 | 0.77 |  | 0.27 | 0.80* | 0.61* | 0.51 |  |  |  |  |  |  |  |  |  |
|  | Cd | Nil | Nil | Nil |  | 0.35 | 0.39 | -0.55 |  |  |  |  |  |  |  |  |  |
|  | Cr | -0.84 | 0.75 | 0.96* | Nil |  | 0.80* | 0.47 |  |  |  |  |  |  |  |  |  |
|  | Pb | -0.67 | 0.84 | 0.99* | Nil | 0.92 |  | 0.19 |  |  |  |  |  |  |  |  |  |
|  | Zn | 0.07 | -0.42 | 0.25 | Nil | 0.27 | 0.11 |  |  |  |  |  |  |  |  |  |  |

Pearson correlation coefficients were shown. Values with \* indicated that significant correlations were found at p=0.05.

Lead pollution in cities was commonly recognized as one of the major pollutants caused by vehicle emissions (Yang *et al.*, 2000). Hong Kong government introduced unleaded petrol (ULP) in 1991 and banned the supply, sale and dispensing of leaded petrol as well as any fuel additives containing Pb in 1999 (Hong Kong Environmental Protection Department, 1999), resulting in a decline of Pb concentration in street dust of Hong Kong from 1300 ± 1400 mg kg^-1^ (Yim and Nau, 1987) to 180 ± 93 mg kg^-1^ (Li *et al.*, 2001).

Atmospheric deposition from nearby regions also represents another important input of heavy metals such as Cr, Cu, Pb and Zn to surface soils. According to a quality monitoring program in China (General Administration of Quality Supervision Inspection and Quarantine of the People’s Republic of China, 2004), only about 70% of the unleaded petrol samples in China was found to comply with the national standard. In some cases, Pb level was exceeded more than 200 times to the standards. Study on atmospheric deposition in the PRD revealed that the deposition of Cr, Cu, Pb and Zn (6.43 ± 3.19, 18.6 ± 7.88, 12.7 ± 6.72 and 104 ± 36.4 mg m^-1^ yr^-1^) was significantly higher when compared with Europe and North America (Wong *et al.*, 2003). Long-range transport of air-borne pollutants or wind-blown contaminated soil particles from Mainland China by the northeast monsoon was reported (Lee and Hills, 2003). Moreover, atmospheric input was reported to be the major contributor of Pb, Cd, As and Hg in agricultural soils in England and Wales (Nicholson *et al.*, 2006), and thus atmospheric deposition, either locally or regionally, may play a significant role for the presence of particle-bound pollutants in soils.

### 3. Comparison of soils cleanup criteria

The soil quality guideline values on the various pollutants investigated imposed by Netherlands (Dutch Guidelines) (Ministry of Housing Spatial Planning and Environment, 2000), Sweden (Soil Remediation Goals) (Swedish Environmental Protection Agency, 2002), England (Kelly Indices) (Contaminated Land Assessment & Remediation Research Centre, 2004) and China (Environmental Quality Standard) (State Environmental Protection Administration of China, 1995) are summarized in Table 3. As Dutch guideline is the most comprehensive and commonly used, comparisons the present findings are made chiefly with the Dutch values. Mercury, Cd and Pb in 138 samples were below the Dutch intervention values, suggesting that the concentrations of these metals in soils were not hazardous to human. In terms of As, Cu, Cr and Zn, most of their levels did not exceed the Dutch intervention values, but there were sporadic soil samples containing levels of As, Cu, Cr and Zn exceeded the intervention values (14, 3, 2, and 8 out of 138 samples correspondingly). Nine out of 10 suburban samples from the northern and northeastern New Territories contained As concentrations greater than the Dutch intervention value. Thus, it is suspected that a large part of northern and northeastern in Hong Kong is contaminated with As at concentration that can impose adverse effects on human health. Soil samples with Zn concentration greater than the intervention value were mainly noted in industrial areas, and this is also true for Cu and Cr. For the scarcity and remoteness of the hotspots (Cr, Cu, and Zn), their potential adverse impacts to the general public were kept to minimum. In England, soil remediation is often required before further development on brownfield soils as they are typically contaminated with high levels of heavy metals due to past industrial activities (French *et al.*, 2006). Although leaching of toxic metals to underground water is not a major concern as Hong Kong relies mainly on river water transported from the mainland as well as rainwater collected locally, the fact that more than 10% of the soil samples were highly contaminated with As warrants further investigation. This is especially true if any of these sites (north and north-east of the New Territories) are used for residential development in the future.

**Table 3.** Soil quality guidelines from Netherlands, Sweden, United Kingdom and China, and their recommended values for Hg, As, Cu, Cd, Cr, Pb and Zn in soils.

| Country of implementation | Quality guidelines | Hg | As | Cu | Cd | Cr | Pb | Zn |
| --- | --- | --- | --- | --- | --- | --- | --- | --- |
|  |  | Concentration in soils (mg/kg) |  |  |  |  |  |  |
| Netherlands | Dutch target value | 0.3 | 29 | 36 | 0.8 | 100 | 85 | 140 |
|  | Dutch intervention value | 10 | 55 | 190 | 12 | 380 | 530 | 720 |
| Sweden | Soil remediation goals for sensitive land use | N.A. | 15 | 100 | 0.4 | 120 | 80 | 350 |
|  | Soil remediation goals for less sensitive land use | N.A. | 40 | 200 | 12 | 250 | 300 | 700 |
| United Kingdom | Kelly | 1 | 30 | N.A. | 1 | 100 | 500 | N.A. |
| China | Environmental quality standard | 0.5 | 30 | 200 | 0.6 | 300 | 300 | 250 |

## 4. Conclusions

In terms of total concentrations of all the metals (metalloids), industrial and highway areas were the most contaminated. This finding is also true for individual elements (As, Cd, Cr, Cu, Pb and Zn) other than Hg. It was found that Hg concentration was the highest in soil collected from agricultural farmland, which could be attributed to the application of Hg-containing agrochemicals. The use of low quality fertilizers is also believed to be the main source of As, Cu and Zn, while substantial contributions of pollutants by exhausts and wearable parts from automobiles are also suspected. Atmospheric deposition from local and nearby regions is also believed to be a major source of edaphic metals in Hong Kong. A reduction of Pb in soils during the past 10 to 15 year was chiefly due to the use of Pb-free petrol. It is expected all the metals (metalloids) would not cause any potential health impacts to the general public, except As, due to its high concentrations in northern and northeastern suburbs in Hong Kong.

## Acknowledgments

The authors are grateful to Mr. Y.Y. Chin from the Leisure and Cultural Services Department (LCSD, HKSAR) for providing technical assistance. Financial support from the Strategic Research Fund, Science Faculty, HKBU and the Area of Excellence (AoE) Scheme (CITYU/AoE/03-04/02) under the University Grants Committee of Hong Kong is gratefully acknowledged.

